# Extended-spectrum β-lactamase genes traverse the *Escherichia coli* populations of ICU patients, staff and environment

**DOI:** 10.1101/2022.12.08.519559

**Authors:** Robert A. Moran, Liu Baomo, Emma L. Doughty, Yingyi Guo, Xiaoliang Ba, Willem van Schaik, Chao Zhuo, Alan McNally

## Abstract

Over a three-month period, we monitored the population of extended-spectrum β-lactam-resistant *Escherichia coli* (ESBL-EC) associated with the patients, staff and environment of an intensive care unit (ICU) in Guangzhou, China. Thirty-four clinical isolates were obtained from the same hospital 12 months later. A total of 165 isolates were characterised and whole-genome sequenced, with 24 isolates subjected to long-read sequencing. The diverse population included representatives of 59 different sequence types (STs). ICU patient and environmental isolates were largely distinct from staff isolates and clinical isolates. We observed five instances of highly similar isolates (0-13 core-gene SNPs) being obtained from different patients or bed unit environments. ESBL resistance in this collection was largely conferred by *bla*_CTX-M_ genes, which were found in 96.4% of all isolates. The contexts of *bla*_CTX-M_ genes were diverse, situated in multiple chromosomal positions and in various plasmids. We identified *bla*_CTX-M_-bearing plasmid lineages that were present in multiple STs across the surveillance, staff and clinical collections. Closer examination of IS*Ecp1*-*bla*_CTX-M_ transposition units shed light on the dynamics of their transmission, with evidence for the acquisition of chromosomal copies of *bla*_CTX-M_ genes from specific plasmid lineages, and for the movement of *bla*_CTX-M-55_ from a ST1193 chromosome to a small mobilisable plasmid. A carbapenem-resistant ST167 strain isolated from a patient that had been treated with meropenem and piperacillin-tazobactam contained seven copies of *bla*_CMY-146_, which appears to have been amplified by IS*1*. Our data revealed limited persistence and movement of ESBL-EC strains in the ICU environment, but we observed circulating plasmid lineages playing an essential and ongoing role in shaping the cephalosporin-resistance landscape in the population examined.

**Impact statement:** ESBL resistance significantly impacts clinical management of *E. coli* infections in hospitals globally. It is important to understand the structures of ESBL-EC populations carried by hospital patients and staff, their capacity to persist in hospital environments, and the dynamics of mobile genes that drive the spread of ESBL resistance. In our three-month study, ESBL-EC strains found in the ICU environment were strongly associated with patient carriage, but distinct from strains found in staff. However, plasmid lineages carrying *bla*_*C*TX-M_ genes were found across the ICU populations and in a collection of clinical isolates obtained one year later. By examining their content and contexts, we have traced the recent histories of chromosomal and plasmid-borne IS*Ecp1-bla*_CTX-M_ transposition units in the ICU population. This allowed us to implicate specific plasmid lineages in the acquisition of chromosomal *bla*_CTX-M_ genes, even when the plasmids were no longer present, and to detect recent transposition of *bla*_CTX-M-55_ from a chromosome to a mobilisable plasmid. Similar high-resolution approaches to the study of mobile genetic elements will be essential if the transmission routes associated with the spread of ESBL resistance are to be understood and subjected to interventions.

**Data summary:** Sequencing reads are available under NCBI BioProject accession PRJNA907549. The 91 complete plasmid sequences generated in this study are in a supplementary file called pDETEC_collection.fa.

## Introduction

*Escherichia coli* occupies a niche in the human gastrointestinal tract that makes it an important vehicle for mobile genes that confer resistance to clinically-relevant antibiotics. Some clones from the vastly diverse *E. coli* population can cause human infections,^1^ so the importance of antibiotic resistance gene carriage by the species is twofold: infections caused by antibiotic-resistant *E. coli* are more difficult to treat, and antibiotic resistance genes carried by human-associated *E. coli* can be transferred to other Gram-negative pathogens. Extended-spectrum β-lactam (ESBL)-resistant *E. coli* (ESBL-EC) usually carry one or more of the various horizontally-acquired β-lactamase (*bla*) genes that can be located in various chromosomal positions or in plasmids. The *bla*_CTX-M_ genes are some of the most clinically important, and have been detected globally in E. coli and other members of the Enterobacterales.^2^ In China, *bla*_CTX-M-55_ has been increasing in prevalence, and in recent years has overtaken *bla*_CTX-M-14_ and *bla*_CTX-M-15_ as the most common ESBL resistance gene seen in ESBL-EC associated with human infections.^3,4^

Dissemination of ESBL resistance genes through global bacterial populations has been facilitated by mobile genetic elements (MGEs).^5^ Plasmid-mediated intercellular transfer plays an obvious role in the horizontal spread of *bla* genes, but the contribution of intracellular transposition is often uncharacterised in population-level studies. Movement of *bla* genes from chromosomal sites to plasmids, or between plasmids, can increase their intercellular transfer potential. Alternatively, transposition from plasmids into chromosomal sites might increase the stability of *bla* genes in new hosts. The insertion sequence IS*Ecp1* is a major driver of intracellular *bla*_CTX-M_ mobility.^5,6^ IS*Ecp1* can mobilise adjacent DNA by recognising alternatives to its right inverted repeat sequence and generating transposition units (TPUs) of various sizes.^5^ Because TPUs can carry sequences from adjacent to their previous insertion site, in some cases it is possible to deduce their recent histories by examining their content.

It is important to understand the diversity and transmission dynamics of both ESBL-EC and *bla* gene-associated MGEs in hospital settings, particularly in intensive care units (ICUs) that host the most vulnerable patients. Although colonisation by antibiotic-resistant *E. coli* has been described as a significant risk for infection in hospitals,^7^ genomic surveillance studies have rarely included ESBL-EC that are not derived from clinical specimens.^8^ Genomic characterisation of ESBL-EC carried asymptomatically by patients or present in hospital environments might provide insights into the dissemination of ESBL resistance. A recent genomic surveillance study of *Klebsiella pneumoniae* in a Chinese ICU highlighted the utility of considering environmental isolates when assessing hospital populations.^9^

Here, we have performed a prospective observational study to examine the ESBL-EC population of an ICU in Guangzhou, China. By sampling the entire ICU patient cohort and the ICU environment weekly, and collecting rectal swabs from ICU staff, we have captured a three-month snapshot of ESBL-EC, ESBL-resistance determinants and their associated MGEs. This allowed us to assess the impact of *E. coli* and MGE transmission on the spread and persistence of ESBL resistance in this setting.

## Materials and Methods

### Ethics

This study was approved by the Medical Ethics Committee of the First Affiliated Hospital of Guangzhou Medical University on May 21^st^, 2018.

### Study design and sampling regimen

This study was conducted in the Internal Medicine ICU of a tertiary care hospital in Guangzhou, China. Sampling occurred weekly over a 13-week period between July and October 2019. Environmental samples were collected from eight single-bed rooms, a six-bed room, and common areas between rooms. A complete list of environmental sampling sites can be found in Table S1. Oral and rectal swabs were obtained from each patient present in the ward on each weekly sampling occasion. Staff rectal and coat swabs were collected on three dates over the course of the study. Swabbing was performed with Copan swabs moistened with Mueller-Hinton broth. Environmental sites were swabbed for 1 minute and transported to the laboratory at room temperature for culturing. Clinical isolates obtained between September and October 2020 were provided by the hospital’s clinical laboratory.

### Bacterial isolation and antibiotic susceptibility testing

Swabs were incubated, shaking, at 37°C in 4 mL of Mueller-Hinton broth until turbidity was observed (usually 16-18 hours, maximum 24 hours). Turbid cultures (50 μL) were spread on CHROMagar ESBL plates and incubated overnight at 37°C. Presumptive *E. coli* colonies were streaked on antibiotic-free Mueller-Hinton agar plates and incubated overnight at 37°C. Single colonies from Mueller-Hinton plates were collected for storage at -80°C, further characterisation and whole-genome sequencing. Species identity was confirmed by MALDI-TOF. Sensitivity to imipenem and meropenem was assessed by broth microdilution according to CLSI guidelines (M100-S26). E. coli ATCC 25922 was used as a quality control strain.

### Plasmid transfer assays

Transfer of the *bla*_CTX-M-55_-bearing plasmid pDETEC16 was assessed by mating host *E. coli* DETEC-P793 with rifampicin-resistant *E. coli* Ec600. DETEC-P793 and Ec600 overnight cultures (100 μL each) were spread on the same Mueller-Hinton agar plate and incubated at 37°C overnight. The resulting lawn was harvested and serially diluted in 0.9% sterile saline. Dilutions were plated on Mueller-Hinton agar containing 20 μg/mL cefepime to select for DETEC-P793, 500 μg/mL rifampicin to select for Ec600, or 20 μg/mL cefepime + 500 μg/mL rifampicin to select for pDETEC16-containing Ec600 transconjugants. Transconjugants were screened for the presence of pDETEC16 and the putative conjugative plasmids pDETEC13, pDETEC14 and pDETEC15 by PCR. Primers and PCR conditions are listed in Table S2.

### Whole genome sequencing and analysis

Genomic DNA was extracted using a Qiagen minikit (Qiagen, Hilden, Germany) in accordance with the manufacturer’s instructions. Whole genome sequencing was performed using both the Illumina HiSeq (Illumina, San Diego, USA) and the Oxford Nanopore GridION (Nanopore, Oxford, UK) platforms (Tianke, Zhejiang, China).

Illumina sequence reads were trimmed and assembled with Shovill v1.1.0 under default settings with a 10x minimum contig coverage (https://github.com/tseemann/shovill). Read quality was determined with FastQC v0.11.8,1 and assemblies were assessed for contamination and completeness using QUAST v5.0.2, CheckM v1.0.13 and ARIBA v2.14.1 with the “Escherichia coli” MLST database. All genomes meeting quality expectations had a total genome size of 4,580,428-5,537,816 bp; N50 ≥ 65,734; GC content of 50.23 – 50.93 %; genome completeness ≥97.46%; ≤2.52% contamination; ≤251 contigs and complete MLST genes without nucleotide heterogeneity.

For hybrid assemblies, Nanopore reads were trimmed with Filtlong v0.2.0 (https://github.com/rrwick/Filtlong) under default settings targeting approximately 100-fold genome coverage. These were assembled with the trimmed Illumina reads using Unicycler v0.4.8^10^ under default settings. For genomes that did not assemble contiguously in this way, Flye v2.7-b1585^11^ was used to assemble long reads first. The resulting Nanopore-only assemblies were input into Unicycler along with short reads under default settings or using bold mode where specified (Table S3). Manual approaches were used to complete some assemblies.

### Genome characterisation

Genomes were initially characterised by using abricate (v0.8.13) to screen with the NCBI AMRFinderPlus and PlasmidFinder databases (both updated 22/09/2021)^12,13^. F-type plasmid replicons were sub-typed using the PubMLST database (https://pubmlst.org/organisms/plasmid-mlst).

Phylogenetic analysis was undertaken for all isolates together, and separately for each ST with more than three isolates. Reference genomes are listed in Table S4. Reference genomes were annotated with Prokka 1.14.0^14^ under default settings. Using Snippy v4.4.5 (https://github.com/tseemann/snippy), isolates from the whole dataset and from each ST were aligned against their appropriate reference genome and a core genome alignment was generated. When more than three isolates were represented in each alignment, recombination was removed using gubbins v2.4.0^15^ with the Fasttree tree builder^16^. SNP-distances were calculated from resulting core-genome alignments with SNP-dists v0.6.3 (https://github.com/tseemann/snp-dists). Phylogenetic trees were constructed with Fasttree v2.1.10 using the nucleotide alignment setting and a general time reversible model^16^.

### Plasmid and translocatable element characterisation

Gene Construction Kit v4.5.1 (Textco Biosoftware, Raleigh, USA) was used to examine and manually annotate plasmid and other mobile DNA sequences.

## Results

### The intensive care unit hosts a diverse E. coli population

Over a three-month period in 2019, 299 samples were collected from ICU patients (59 ESBL-EC-positive; 19.7%), 82 from ICU staff (38 ESBL-EC-positive; 46.3%) and 2967 from the ICU environment (110 ESBL-EC-positive; 3.7%). A total of 131 ESBL-EC isolates were sequenced (Figure 1, Table S5). Sequenced ICU surveillance isolates were derived from patient oral swabs (10 isolates) and rectal swabs (38 isolates), the ICU environment (47 isolates), and from staff rectal swabs (36 isolates). Sinks were the most common sources of environmental isolates (32 of 47 isolates, 68.1%), which were obtained from sink countertops (12 isolates), overflows (9 isolates) drains (8 isolates), taps (2 isolates) or water (1 isolate). The remaining environmental isolates were found on bed unit or equipment surfaces, including those of bed remotes (5 isolates) and bed curtains (1 isolate), a locker (1 isolate), ventilators (2 isolates), a nebuliser (1 isolate) and drip stands (2 isolates). One isolate was collected from a door handle, one from a cleaning cart and one from a doctor’s coat. A further 34 ESBL-EC isolates were obtained from clinical samples taken from patients throughout the hospital over a two-month period in 2020.

**Figure 1:**
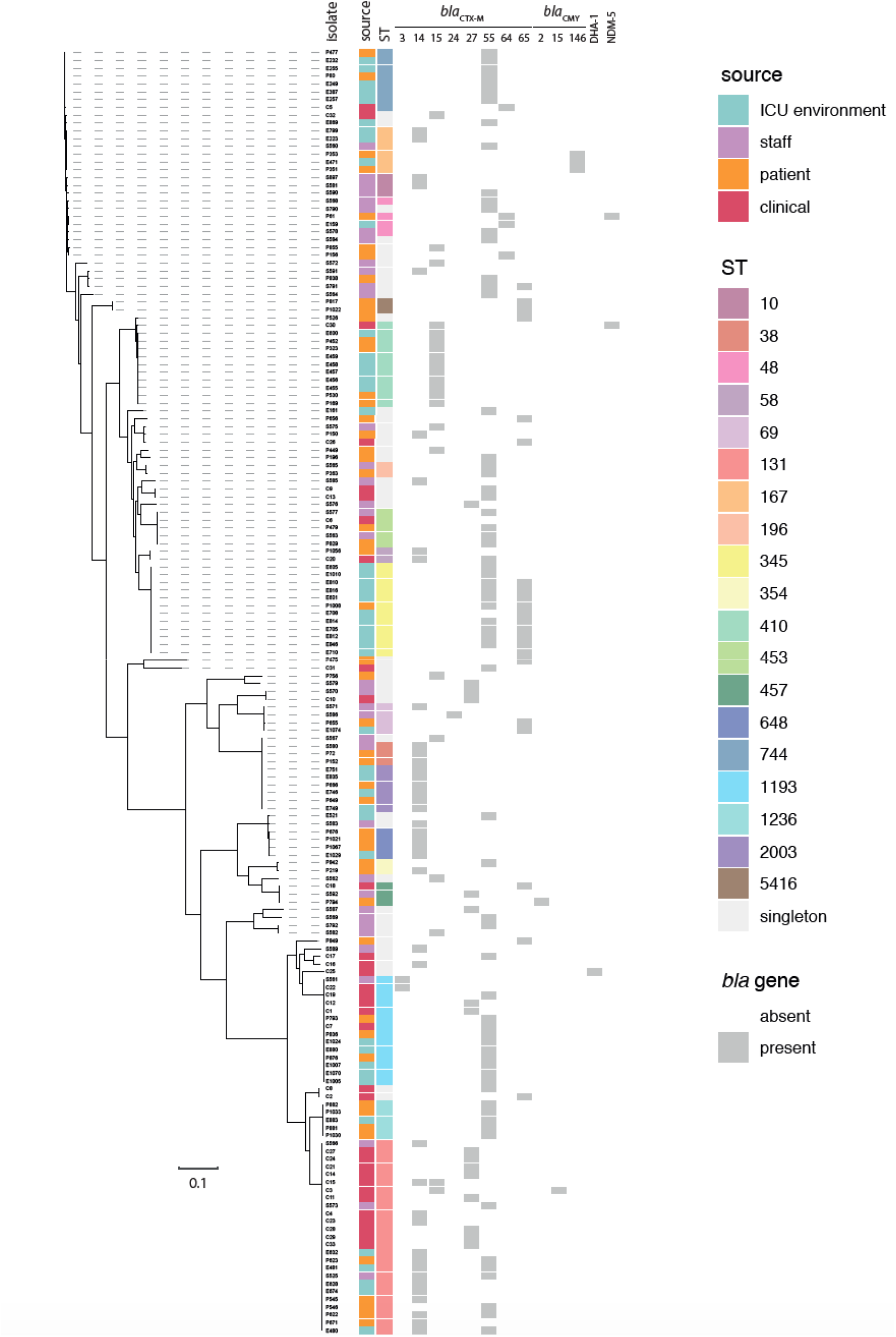
ESBL-EC collection phylogeny. Core-gene phylogeny of the ESBL-EC collection assembled in this study. Isolate names are labelled to the right of dashed lines that indicate their positions in the phylogeny. To the right of the phylogeny, sources of isolation, sequence type (ST) designations and the presence or absence of *bla* genes are indicated by colours as outlined in the key. High-resolution figure included with supplementary materials.

Multi-locus sequence typing revealed 50 different sequence types (STs) in the ICU surveillance collection (131 isolates) and 17 in the clinical collection (34 isolates). One ICU surveillance isolate and one clinical isolate were novel types, which were submitted to Enterobase and assigned ST12546 and ST12742. Of the 59 STs in the entire collection, 36 were only represented by single isolates and 19 were represented by between two and eight isolates. The most prevalent STs in the collection were ST131 (25 isolates), ST1193 (14), ST345 (11) and ST410 (11). Eight of the 17 STs in the clinical collection were also present in the ICU surveillance collection, namely ST131, ST1193, ST410, ST744, ST453, ST58, ST393 and ST457.

### Environmental ESBL-EC isolates were strongly associated with patient carriage

Visualising the distribution of ESBL-EC isolates, patients and bed units over the course of the surveillance study revealed patterns of ST, patient and environmental associations in the ICU (Figure 2). ESBL-EC was isolated from a patient or their bed unit on 60 sampling occasions that involved 30 different patients. On 32 of these occasions, isolates were derived from only the patient; on 15 occasions from only the environment; and on 13 occasions from both the patient and their bed unit environment. On nine of the 13 occasions when ESBL-EC was isolated from both the patient and their bed unit environment, the environmental and patient isolates were the same ST. Of the 15 occasions on which ESBL-EC was isolated from a bed unit environment but not its resident patient, in nine the environmental isolate’s ST was the same as isolates that had been collected from that patient in the week(s) prior. Thus, of 60 sampling occasions where ESBL-EC was isolated from occupied bed units, 50 occasions (83.3%) involved STs obtained directly from occupying patients at the time of sampling or on previous sampling occasions.

**Figure 2:**
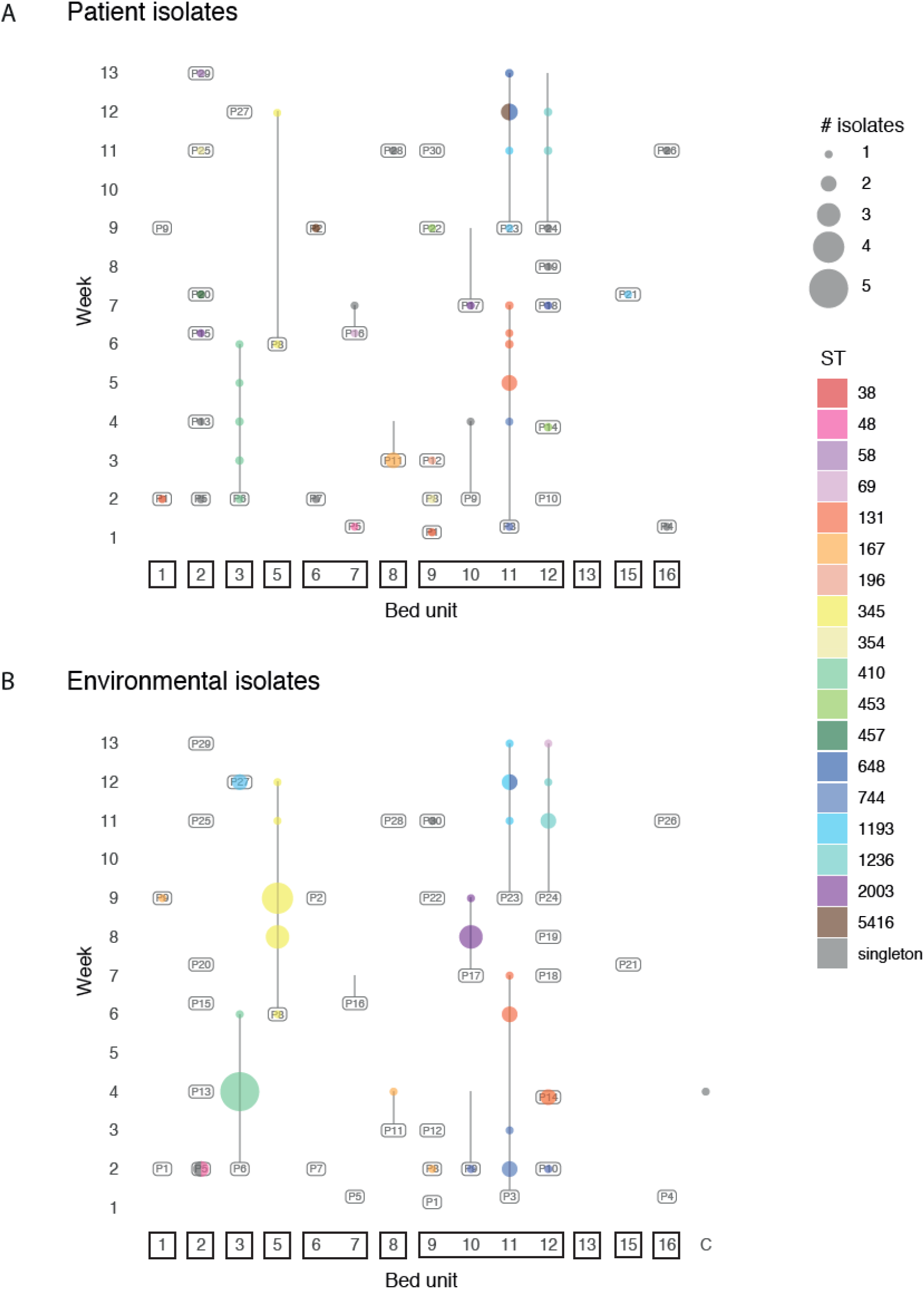
Distribution of ESBL-EC STs in the ICU surveillance study. Bubble plot showing the distributions of **A)** patient-derived and **B)** ICU environment-derived ESBL-EC isolates over the course of the ICU surveillance study. The locations in which STs were isolated are indicated by coloured bubbles, with the sizes of bubbles indicative of the number of isolates obtained. C = common areas outside bed units.

In all 11 cases where a patient and their bed unit were sampled longitudinally, at least one ST was isolated from patient or bed unit on multiple sampling occasions. ESBL-EC isolates associated with a patient and their bed unit were usually a single ST throughout the patient’s ICU stay. Only five patients (P3, P8, P16, P23, P24) were associated with carriage of multiple STs, with those different STs isolated on separate sampling occasions (Figure 2A). Two patients (P1, P8) moved between bed units during the study, and yielded ESBL-EC from oral or rectal swabs in both locations. In both cases the same ST was isolated in both locations (Figure 2A).

### Evidence for strain persistence and dissemination in ICU environments

Across the ICU surveillance and clinical collections, 19 STs were associated with multiple environments, patients or staff members. To determine whether isolates of the same ST were closely related and might be derived from a single introduction to the ICU, we evaluated core-genome SNP (cgSNP) distances as well as plasmid replicon and antibiotic resistance gene content. Where available, whole genome sequences were also compared to confirm the relationships between closely-related isolates. Distances between isolates of the same ST ranged from 0 to 20,795 cgSNPs (median= 311 SNPs; IQR = 114 - 4536 SNPs; Table S6). The median maximum cgSNP distance between isolates of the same ST associated with a single patient was 3 cgSNPs (IQR = 1 - 9 cgSNPs), though up to 99 cgSNPs were found between ST345 isolates associated with P8 (Table S7). Cases where closely related isolates of the same ST were present in multiple bed units are outlined below.

ST744 isolates were obtained from adjacent bed units 10, 11 and 12 between weeks 1 and 4 (Figure 2). ST744 first appeared in BU11 in week 1, isolated from a P3 rectal swab. It was then isolated from the BU11 environment in weeks 2 and 3 before it was isolated from another P3 rectal swab in week 4. Isolates from P3 and BU11 differed by a maximum of three cgSNPs. In week 2, ST744 isolates were also obtained from the environments of BU10 and BU12, which are in the same room as BU11 (Figure 2B). The BU10 and BU12 isolates differed from the BU11 isolates by 1-2 cgSNPs and 9-13 cgSNPs, respectively. All ST744 isolates carried the same ARGs and plasmid replicons.

The ST744 strain in P3 appears to have been displaced by a ST131 strain over P3’s time in the ICU. From week 5 until their discharge from the ICU after sampling in week 7, P3 yielded ST131 isolates from oral and rectal swabs (Figure 2A). ST131 isolates were also obtained from the BU11 environment in weeks 6 and 7 (Figure 2B). The P3 ST131 isolates differed by 1-8 cgSNPs from two isolates obtained from equipment in the adjacent BU12 a week earlier. Complete genomes were obtained for DETEC-E480, isolated from the BU12 environment in week 4, and DETEC-P622, isolated from a P3 rectal swab in week 6. Both genomes contain six plasmids, five of which are identical (Figure S1). The sixth plasmid in each genome is an FII-33:N cointegrate that contains multiple antibiotic resistance genes. The FII-33:N plasmid in DETEC-E480, pDETEC56, is 103,838 bp and pDETEC60 in DETEC-P622 is 82,673 bp. The difference in size is accounted for by an IS*26*-mediated deletion event which has removed the *fosA3, sul2, strAB, tet*(A) and *floR* genes from pDETAB60, leaving only *rmtB, bla*_TEM_ and *bla*_CTX-M-55_ (Figure S1). One ST131 isolate obtained from P3 in week 5, and all ST131 isolates from P3 or BU11 after week 6 did not contain FII-33 or N replicons, or the resistance genes associated with the FII-33:N cointegrate, suggesting that this plasmid had been lost. A further ST131 isolate that differed from those in P3/BU11 by 0-8 cgSNPs and contained the FII-33:N replicons and associated ARGs was isolated from a doctor’s coat in week 8.

After P3 had been discharged, P23 occupied BU11 from week 9 to the end of the study in week 13. Over this period, 11 ST1193 isolates were isolated from P23 and the BU11 environment, including from the sink. These isolates differed by 0-8 cgSNPs. In week 12, two ST1193 isolates were obtained from the sink in BU3 (Figure 2B). The BU3 sink isolates differed from the P23/BU11 ST1193 isolates by 0-7 cgSNPs, and all BU11/BU3 isolates carried the same ARGs and plasmid replicons. Complete genomes were obtained for the ST1193 isolates DETEC-P836 from a P23 rectal swab, DETEC-E1005 from the BU3 sink and DETEC-E1070 from the BU11 sink. All three genomes contain the identical plasmids pDETEC3 and pDETEC4.

Other examples of closely related isolates from different patients or ICU environments include an ST5416 isolated in week 12 from P23 from an oral swab. This ST had only previously been isolated from P2 in bed unit 6, in week 9, which was the same week P23 was admitted to the ICU. The ST5416 isolates differed by 1 cgSNP and contained the same ARGs and plasmid replicons. ST167 isolates with 0 cgSNPs were found seven weeks apart, from the bed curtain of BU9 in week 2 and from the sink overflow of BU1 in week 9, associated with P8 and P9, respectively (Figure 2B). P8 had occupied BU9 in week 2, at which point they were adjacent to P9, who was in BU10 from week 2 to week 4. After a four-week absence from the ICU, P9 was in BU1 when ST167 was isolated from its sink overflow. ST174 isolates that differed by 1 cgSNP were obtained from rectal swabs from two different staff members, but they carried different *bla*_CTX-M_ genes. ST393 isolates from a staff rectal swab and a clinical specimen from 2020 carried *bla*_CTX-M-27_ had identical core genomes (0 cgSNPs). These appeared to represent the only ESBL-EC strain found in both the clinical and ICU surveillance collections.

### Diverse ESBL resistance determinants were found in diverse genetic contexts

CTX-M-type β-lactamases were the dominant ESBL resistance determinants in this collection, with one or more *bla*_CTX-M_ genes found in 159 of the 165 isolates (96.4%). Most isolates (143/159; 89.9%) contained a single *bla*_CTX-M_ gene, while 16 contained two different *bla*_CTX-M_ genes (Figure 1). The *bla*_CTX-M-55_ gene was the most common in the collection (67 isolates; 27 STs), followed by *bla*_CTX-M-14_ (41 isolates; 15 STs), *bla*_CTX-M-15_ (23 isolates; 7 STs), bla_CTX-M-65_ (22 isolates; 11 STs), *bla*_CTX-M-27_ (15 isolates; 7 STs), *bla*_CTX-M-3_ (2 isolates; both ST1193) and *bla*_CTX-M-24_ (1 isolate; ST69). Amongst the isolates that carried two *bla*_CTX-M_ genes, nine had bla_CTX-M-55_ with *bla*_CTX-M-65_, six had *bla*_CTX-M-55_ with *bla*_CTX-M-14_ and one had *bla*_CTX-M-15_ with *bla*_CTX-M-14_. Of the six that lacked *bla*_CTX-M_ genes, three ST167 isolates carried *bla*_CMY-146_, a ST457 isolate carried *bla*_CMY-2_, a ST706 isolate carried bla_DHA-1_ and a ST453 isolate carried only *bla*_TEM_ (Figure 1).

We determined the context of *bla*_CTX-M_ genes in 93 of the 165 isolates in the collection, by examining complete genomes (23 isolates) or *bla*_CTX-M_-containing contigs in draft genomes (70 isolates). In the remaining cases *bla*_CTX-M_ genes were found in contigs that only included mobile element sequences and therefore did not contain sufficient information to reliably determine their locations. Of the 93 instances where the locations of *bla*_CTX-M_ genes were determined, 44 were in chromosomes and 50 in plasmids (one isolate carried copies of *bla*_CTX-M-55_ n its chromosome and in a plasmid). In 55 cases *bla*_CTX-M_ genes were located in complete IS*Ecp1* TPUs for which boundary sequences could be determined. The sizes of these TPUs ranged from 2,841 bp to 18,201 bp (Table S5).

The 55 complete IS*Ecp1-bla*_CTX-M_ TPUs were inserted in 18 different positions in chromosomes and seven in plasmids (Table S5). All complete TPUs were flanked by 5 bp target site duplications (TSDs). Three TPU-insertion position combinations were seen in multiple STs. A 2,845 bp chromosomal unit was flanked by the TSD TGTTT in position in five ST1236 isolates, and one isolate each of ST1485 and 3941. The other combinations found in multiple STs were associated with I-complex plasmids: a 2,971 bp unit in an I1 plasmid was in five STs and a 3,060 bp unit in a Z plasmid was in three STs. This suggested that *bla*_CTX-M_-bearing I1 and Z-type plasmid lineages might be circulating in this *E. coli* population.

### I-complex plasmid lineages found in multiple STs

To investigate the possibility that the same I1 and Z plasmid lineages were present in multiple STs in this *E. coli* population, we compared complete plasmid sequences to one another and to contigs from draft genomes that represent incomplete plasmid sequences.

I1 plasmids containing a 2,971 bp IS*Ecp1-bla*_CTX-M-55_ TPU flanked by the TSD TACTT were found in six isolates in this collection: one ST1011 and two ST1193 isolates from clinical specimens, and one isolate each of ST93, ST167 and ST196 from ICU staff rectal swabs. The backbones of these plasmids were typical representatives of the I1 type (Figure 3A), containing shufflons and complete transfer regions like those of the reference plasmid R64.^17^ Based on the presence of two recombinant patches in their backbones, the I1 plasmids in this collection were divided into two sub-lineages, represented by pDETEC69 and pDETEC73 in Figure 3A. The plasmids from ST93 and ST167 isolates belonged to the pDETEC69 sub-lineage, while those from ST196, ST1011 and ST1193 isolates belonged to the pDETEC73 sub-lineage.

**Figure 3:**
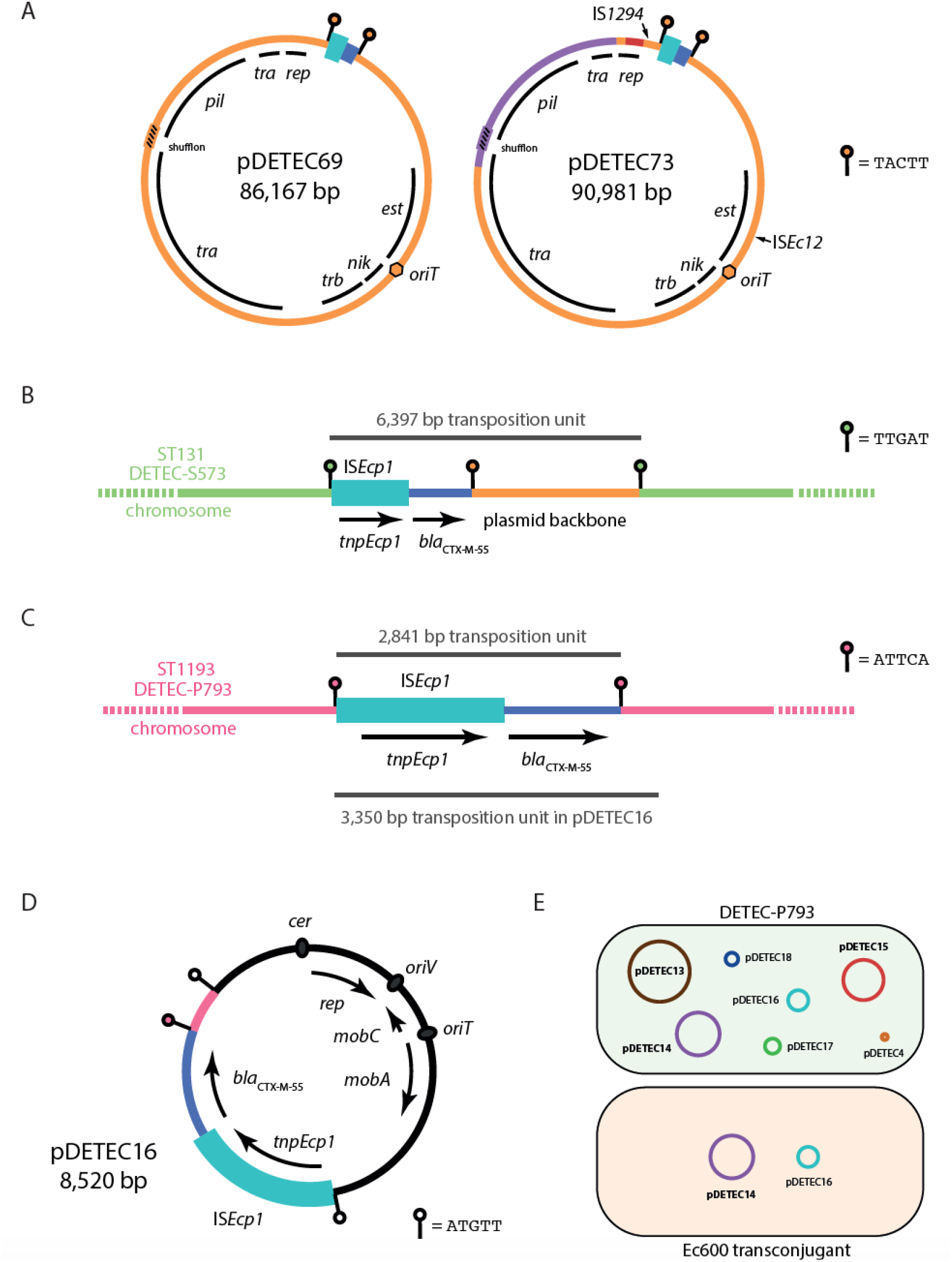
Plasmids, transposition units and *bla*_CTX-M_ movement. **A)** Circular maps of the I1 plasmids pDETEC69 and pDETEC73. The extents of replication (*rep*), transfer (*tra, trb*), thin pilus biogenesis (*pil*) and establishment (*est*) regions are shown. IS*Ecp1-bla*_CTX-M_ TPUs are shown as cyan/blue boxes flanked by lollipops that indicate the position and sequence of target site duplications. Purple and maroon segments in pDETEC73 represent recombinant sequences. **B)** IS*Ecp1-bla*_CTX-M-55_ TPU in the chromosome of ST131 isolate DETEC-S573. **C)** IS*Ecp1*-*bla*_CTX-M-55_ TPU in the chromosome of ST1193 isolate DETEC-P793. **D)** Small *bla*_CTX-M-55_-bearing plasmid pDETEC16. **E)** Co-transfer of pDETEC16 and pDETEC14. Shaded cells represent DETEC-P793 and a transconjugant derived from mating DETEC-P793 with *E. coli* Ec600. The plasmids in each host are shown as labelled circles. Parts A to D of this figure are drawn to different scales, though IS*Ecp1* (1,656 bp) is shown in each, and the sizes of TPUs in parts B and C are indicated.

Z plasmids containing a 3,050 bp IS*Ecp1-bla*_CTX-M-14_ TPU flanked by the TSD GCGGA were found in four isolates in this collection: a ST131 isolate from a clinical specimen, a ST58 isolate from an ICU patient rectal swab, and ST95 and ST131 isolates from ICU staff rectal swabs. Similar to the situation seen amongst I1 plasmids, the Z plasmids could be divided into sub-lineages on the basis of backbone recombination patches. Plasmids from the patient ST58 (pDETEC82) and staff ST131 (pDETEC79) isolates belonged to the same sub-lineage. Apart from rearrangements in the shufflon region (which was also interrupted by IS*1* in pDETEC82), pDETEC79 and pDETEC82 were almost identical (99.98% nucleotide identity across 85,765 bp compared).

Both of the signature TPU-backbone junction sequences from the I1 and Z plasmids described above were found in multiple plasmids in GenBank, indicating that these lineages are present in wider enterobacterial populations. Plasmids bearing the I1 plasmid TACTT-flanked IS*Ecp1-bla*_CTX-M-55_ insertion (n = 25) have been seen in *E. coli, Shigella sonnei, Salmonella Typhimurium, Klebsiella pneumoniae* and *Enterobacter hormachei* that were isolated from human faeces, clinical isolates, animals and wastewater in China (n = 18), Japan (n = 3), Kazakhstan, Belgium, Switzerland and the UK (n = 1 each) (Table S8). Plasmids containing the Z-plasmid GCGGA-flanked IS*Ecp1-bla*_CTX-M-14_ insertion (n = 44) have been carried by *E. coli, K. pneumoniae, Salmonella* and *Shigella* isolated from multiple countries in Asia and Europe, as well as in Australia and the USA (Table S9).

### Evidence linking chromosomal *bla*_CTX-M_ genes to specific plasmid lineages

To investigate the dynamics of their inter-host and inter-molecular transmission, we examined the contents of complete chromosomal IS*Ecp1-bla*_CTX-M-55_ TPUs. In six cases we were able to definitively identify the plasmid lineages that chromosomal insertions were derived from. In the ST131 isolate DETEC-S573 from an ICU staff rectal swab, *bla*_CTX-M-55_ is located in a 6,397 bp IS*Ecp1* TPU inserted in the chromosome and flanked by the TSD TTGAT (Figure 3B). The 6,397 bp TPU includes 3,426 bp of I1 plasmid backbone from immediately adjacent to the 2,971 bp TPU described above, including one copy of the associated TSD sequence TACTT. Thus, we conclude that the 6,397 bp TPU in this ST131 chromosome is derived from the I1 plasmid lineage present in multiple STs in this ESBL-EC population (Figure 3B). As DETEC-S573 does not contain an I1 plasmid, the plasmid must have been lost after delivering the *bla*_CTX-M-55_ gene. Similarly, we determined that a HI2 plasmid lineage (GenBank accession MT773678) was the source of the 18,201 bp chromosomal IS*Ecp1-bla*_CTX-M-64_ TPU in two ST48 isolates, a second HI2 plasmid lineage (AP023198) the source of the 3,050 bp chromosomal IS*Ecp1-bla*_CTX-M-55_ TPU in a ST12742 isolate, an I2 plasmid (LR890295) the source of the 5,800 bp chromosomal IS*Ecp1-bla*_CTX-M-55_ TPU in a ST617 isolate, and an I-complex plasmid lineage not represented in this collection or in GenBank was the source of the 3,445 bp chromosomal IS*Ecp1-bla*_CTX-M-14_ TPU in a ST345 isolate (Table S5). The FII-2 plasmid lineage represented by pHK01 (HM355591), which is present in clinical ST12 isolate DETEC-C16 from this collection, was the source of the 4,477 bp chromosomal IS*Ecp1-bla*_CTX-M-14_ TPU in two ICU patient ST38 isolates (Table S5).

### Chromosome-to-plasmid transposition of *bla*_CTX-M-55_ in ST1193

The complete genome of ST1193 patient rectal isolate DETEC-P793 contained two copies of *bla*_CTX-M-55_, one in the chromosome and one in a small plasmid. The chromosomal copy is in a 2,841 IS*Ecp1* TPU (Figure 3C). The second copy is in the 8,520 bp ColE2-like plasmid pDETEC16 (Figure 3D). The IS*Ecp1-bla*_CTX-M-55_ TPU in pDETEC16 is 3,350 bp and flanked by the 5 bp target site duplication ATGTT (Figure 3D). The final 509 bp of the TPU are identical to the sequence adjacent to the DETEC-P793 chromosomal TPU (Figure 3D). This indicates that the TPU in pDETEC16 was acquired from its host’s chromosome. pDETEC16 has a ColE2-like backbone that contains a putative origin-of-tranfer (*oriT*) and MOB_Q4_-type mobilisation determinants (Figure 3D).

Three of the seven plasmids carried by DETEC-P793 contain complete transfer regions (Figure 3E). We mated DETEC-P793 with *E. coli* Ec600 in order to determine whether any of the large plasmids in DETEC-P793 could mobilise pDETEC16. Transconjugants were obtained at a mean frequency of 8.55 × 10^−6^ per donor. Five transconjugants were screened for the presence of pDETEC16 and all three putative conjugative plasmids by PCR. The I1 plasmid pDETEC14 was detected along with pDETEC16 in all transconjugants, while pDETEC13 and pDETEC15 were not detected in any (Figure 3E). This demonstrated that pDETEC14 had mobilised pDETEC16 in the laboratory. Mobilisation of pDETEC16 by an I1 plasmid is consistent with previous studies that have shown that MOB_Q4_-type plasmids can be mobilised by I-complex plasmids.^18^

### Carbapenem resistance associated with IS*1*-mediated amplification of *bla*_CMY-167_

All isolates were tested for susceptibility to meropenem, and just four exhibited resistance. Carbapenem resistance in the ST410 clinical isolate DETEC-C6 and the ST48 patient rectal swab isolate DETEC-P61 could be explained by the presence of the *bla*_NDM-5_ metallo-ß-lactamase gene. The two remaining meropenem-resistant isolates were ST167 and contained *bla*_CMY-167_, which is not expected to confer resistance to carbapenems. DETEC-P351 was isolated from a P11 rectal swab in week 3 and DETEC-E471 from P11’s bed unit environment a week later (Figure 2). The complete genome of DETEC-P351 contains seven copies of *bla*_CMY-167_. Six copies are in the 77,960 bp I1 plasmid pDETEC6, and the seventh is in the chromosome (Figure 4A). The copies of *bla*_CMY-167_ in pDETEC6 are interspersed with copies of IS*1*, in a configuration that resembles structures produced by IS*26*.^19^ Consistent with amplification of *bla*_CMY-167_ by IS*1* in the I1 plasmid context, we found a putatively ancestral I1 plasmid in GenBank (pECY56; accession KU043116) that contains just a single copy of *bla*_CMY_, with flanking sequences identical to those in pDETEC6 (Figure 4B).

**Figure 4:**
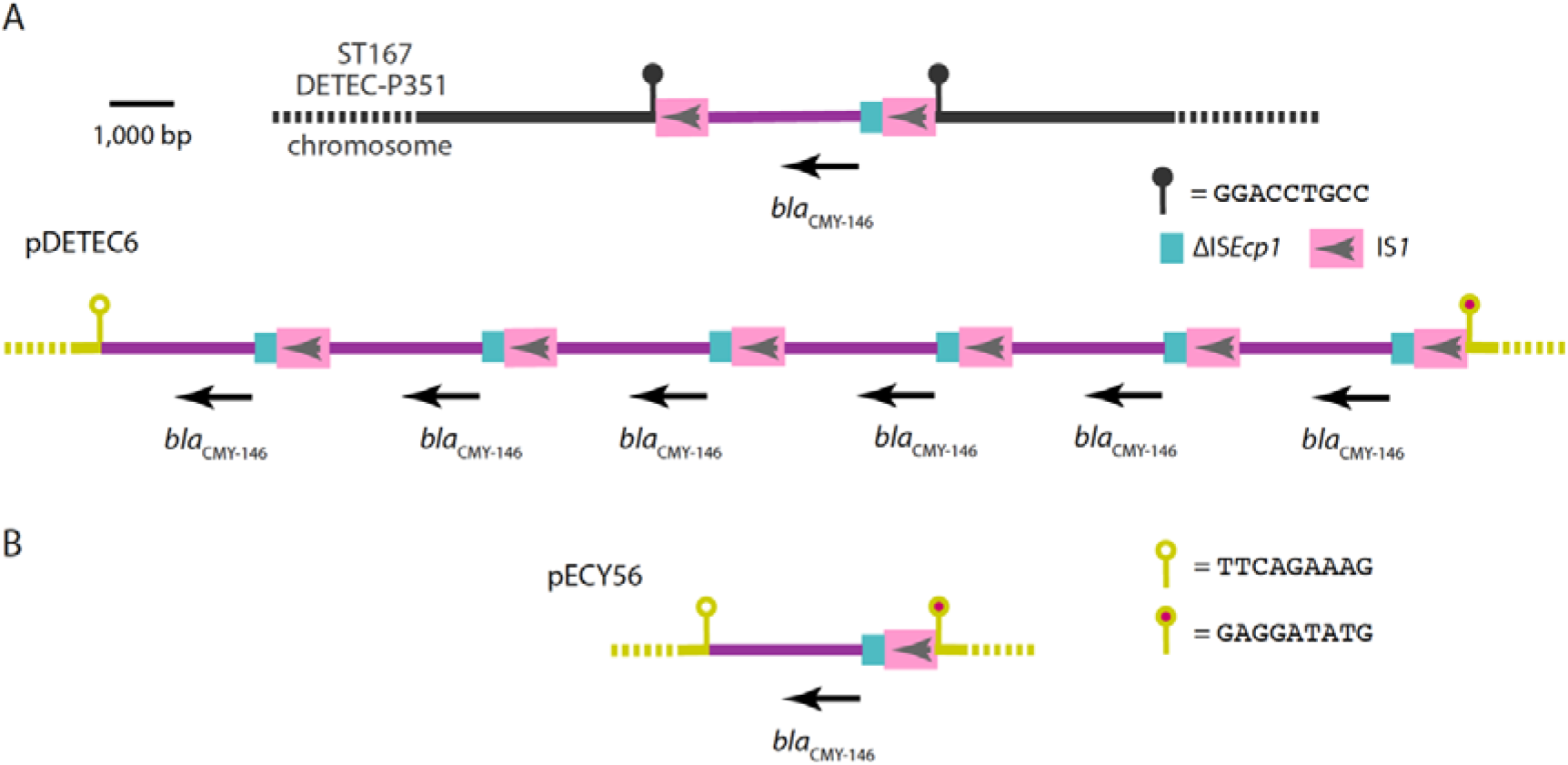
Amplification of *bla*_CMY-146_ in *E. coli* ST167. Scaled diagrams showing **A)** Contexts of *bla*_CMY-146_ in DETEC-P351, and **B)** The context of *bla*_CMY_ in pECY56. IS*1* are shown as pink boxes with arrows indicating the orientation of their transposase genes. Fragments of IS*Ecp1* are shown as cyan boxes, and the amplified sequence containing *bla*_CMY_ as purple lines. The DETEC-P351 chromosome is shown as a black line, and the pDETEC6/pECY56 backbone as a staggered grey line. Colour-filled lollipops indicate the positions of the sequences shown.

The chromosomal *bla*_CMY-167_ gene in DETEC-P351 lies between two copies of IS*1* in what appears to be a 4,266 bp compound transposon flanked by the 9 bp TSD GGACCTGCC (Figure 4A). The 2,730 bp passenger sequence between the copies of IS*1* is identical to the amplified segment in pDETEC6. The chromosomal copy of *bla*_CMY-167_ is therefore likely to have been acquired from pDETEC6.

### Internationally-distributed multi-drug resistance plasmid lineages in the ICU

We have generated 91 complete plasmid sequences as part of this study (Table S10). These represent a diverse spectrum of plasmid types, including commonly-described large plasmids, phage-plasmids, and small plasmids that utilise rolling-circle replication or theta replication with RNA (θ-RNA) or protein (θ-Rep) initiators. Forty of the complete plasmids contain one or more antibiotic resistance genes. Most ARG-containing plasmids were F-types (n=24) or I-complex (n=10), with the remainder X-types (n=2), a phage-plasmid, a H-type plasmid, a θ-RNA plasmid and a θ-Rep plasmid (n=1 each).

Amongst the F-type plasmids we found examples of well-characterised internationally-distributed lineages. Four complete plasmids contained FII-33 replicons, and when we examined draft genomes we found FII-33 replicons in a further 22 isolates. We have recently described the diversity and evolution of the FII-33 plasmid lineage, which is endemic in China, internationally-disseminated and strongly associated with multi-drug resistance in *E. coli* and *K. pneumoniae*.^20^ Five complete plasmids were members of F-type ColV/ColBM lineages that carry colicin and virulence genes, and a further 27 draft genomes contain all or part of the *cvaC* colicin V gene. The virulence-associated genes in ColV/ColBM plasmids include those for siderophores such as aerobactin and salmochelin, which are thought to contribute to extra-intestinal virulence in *E. coli*.^21,22^ Acquisition of these plasmids has played an important role in the evolution of some pathogenic *E. coli* lineages, and they have been associated with pandemic lineages such as ST131, ST95 and ST58.^23,24^ ColV and ColBM plasmid lineages are known to have acquired antibiotic resistance determinants,^24^ and all five complete examples in this collection contained multiple resistance genes in complex resistance regions.

## Discussion

This study has provided a high-resolution three-month snapshot of an ICU’s ESBL-resistant *E. coli*. The population was diverse, with strains carried by staff largely distinct from those found in patients and the ICU environment. ESBL resistance determinants were also diverse, and although *bla*_CTX-M-55_ and *bla*_CTX-M-14_ dominated, they were found in various contexts in plasmids and chromosomes. Some *bla*_CTX-M_-bearing plasmid lineages were found across the disparate E. coli populations, or were shown to have introduced *bla*_CTX-M_ genes that transposed into host chromosomes as passengers in IS*Ecp1* TPUs. Our close examination of IS*Ecp1* TPUs also allowed us to detect the movement of *bla*_CTX-M-55_ from a chromosomal site to a mobilisable plasmid in a ST1193 strain (Figure 3C-E).

There was a strong relationship between isolates found in ICU patients and those found in their bed unit environments. However, we observed limited strain persistence in the ICU environment. Although instances of highly-similar isolates being found in multiple bed unit environments were rare, we observed more involving units in the six-bed room (BU9-12; ST744, ST131 and ST1193) than we did other rooms (Figure 2). This suggests that ESBL-EC transmission is more likely to occur in multi-bed ICU rooms. Of the three instances where highly-similar isolates were found in bed units in different rooms, two (involving ST1193 and ST167) were associated with sinks. Hospital sinks have been shown in other studies to be important reservoirs of antibiotic-resistant pathogens,^25,26^ and to contribute to transmission via plumbing in model systems.^27^

Although there was little crossover at strain level between the ICU and clinical collections, some *bla*_CTX-M_-bearing plasmid lineages were represented in both, as well as in multiple STs within the ICU surveillance collection. I-complex plasmids (I1 and Z types) were particularly prominent here. The association of *bla*_CTX-M_ genes with I-complex plasmids has been noted internationally, and the existence of multiple internationally-disseminated lineages ^28^ suggests that the confluence of these elements has proven successful on many occasions. However, where and under which conditions these and other plasmids are transferring in bacterial populations remain open questions. We did not find evidence here for horizontal transfer of plasmids in the ICU, though our examination of only a single ESBL-EC colony per sample precluded this.

The diversity of the ICU ESBL-EC population, and its strong association with patient or staff carriage, appears to suggest that new ESBL-EC strains are introduced to the ICU regularly. The 46.3% ESBL-EC carriage rate observed in staff here is indicative of a high community carriage rate, as the ICU staff are healthy adults residing in Guangzhou. This highlights the importance of genomic studies targeting community commensal *E. coli* populations,^29^ which might reveal links to the strains and plasmids that are ultimately associated with hospital infections.

A concerning finding here was the presence of multiple copies of *bla*_CMY-146_ in a carbapenem-resistant ST167 strain that lacked carbapenemase genes (Figure 4). This appears to be another example where IS-mediated amplification of a β-lactam^30–32^ or aminoglycoside^33^ resistance gene has yielded an unexpected phenotype. In previous cases IS*26* has been involved in gene amplification, but here IS*1* was implicated. As IS*1* is not part of the IS*26* family of elements, for which study of transposition mechanisms has provided an explanation for observed structures,^34^ similar molecular examinations of IS*1* transposition are required. More generally, the modulation of clinically-relevant β-lactam resistance phenotypes by IS-mediated gene duplications requires further investigation.

## Conclusions

The patients, staff and environment of this ICU hosted a diverse ESBL-EC population over our three month-study period. Our data suggest that strains are being introduced to the ICU regularly, likely in association with patients, but that these strains do not persist for extensive periods in ICU environments. Plasmid and IS*Ecp1*-mediated transmission of *bla*_CTX-M_ genes play major roles in the ongoing spread of ESBL resistance in *E. coli* populations that can enter hospitals.

