## Supplementary files for "Extended-spectrum β-lactamase genes traverse the *Escherichia coli* populations of ICU patients, staff and environment"

### Supplementary Tables and Figure

**Table S1:** Environmental sampling locations.

| Bed units | Communal areas |
|---|---|
| bed rail and regulator | computer: keyboard, mouse |
| ventilator (humidifier, display screen, connector) | phone |
| ECG monitor | barcode scanner |
| micropump | blood sample rack |
| switch button | medical record rack |
| nebuliser | sinks and pipe |
| stethoscope, flashlight | water dispenser |
| inner wall of hanging tower and drawer handle | cabinet and table |
| computer: keyboard, mouse | switch buttons |
| lockers | fibrobronchoscope |
| treatment vehicle | refrigerator |
| air disinfector | rescue vehicle |
| door | defibrillator |
| bed curtain | blood gas analyser |
| sink: tap | ultraviolet steriliser |
| sink: faucet surface | blood filter |
| sink: pool table | ECG machine |
| sink: drain | non-invasive ventilator |
| sink: water pipe | laryngoscope |
| | oxygen tank |
| | ice machine |
| | cleaning cart |
| | mop handle |
| | door handles |

**Table S2:** Detection of DETEC-P793 plasmids by PCR.

| Primer name | Sequence (5'-3') | Product size (bp) | Plasmid detected |
|---|---|---|---|
| repDETEC13-FW | CCTGTTGCTTTGTTGAGGCT | 592 | pDETEC13 |
| repDETEC13-RV | AGATTGGTGCGGGGTCTTTA | | |
| RH1651 | TTAGCACCCGAAGAGCAGAT | 948 | pDETEC14 |
| RH1652 | CTGACGCAACTCCCTGATG | | |
| FIA-FW | CCATGCTGGTTCTAGAGAAGGTG | 462 | pDETEC15 |
| FIA-RV | GTATATCCTTACTGGCTTCCGCAG | | |
| repDETEC16-FW | CACGGATGATCTCGCTTTCG | 760 | pDETEC16 |
| repDETEC16-RV | GCGCTTAGACTTAGAACCGC | | |

All PCRs were performed with an annealing temperature of 60°C.

Primers repDETEC13-FW/RV and repDETEC16-FW/RV were designed for this study.

Primers RH1651/1652 have been published in Moran *et al* 2015 (PMID 25819400) and FIA-FW/RV have been published in Carattoli *et al* 2005 (PMID 15935499).

**Table S3:** Overview of complete[1] genomes generated in this study.

| Isolate[2] | Date | Source | Assembly[3] | ST | #ARGs[4] | #plasmids |
|---|---|---|---|---|---|---|
| DETEC-P61 | 01/08/2019 | rectal swab | FUM | 48 | 22 | 6 |
| DETEC-P1056 | 22/10/2019 | rectal swab | FU | 58 | 2 | 6 |
| DETEC-S586 | 01/09/2019 | rectal swab | U | 69 | 5 | 10 |
| DETEC-S589 | 01/09/2019 | rectal swab | U | 95 | 3 | 2 |
| DETEC-E480 | 19/08/2019 | switch button | FU | 131 | 14 | 6 |
| DETEC-P622 | 03/09/2019 | rectal swab | FU | 131 | 9 | 6 |
| DETEC-S566 | 01/09/2019 | rectal swab | FU | 131 | 3 | 2 |
| DETEC-P351 | 13/08/2019 | rectal swab | U | 167 | 7 | 6 |
| DETEC-E223 | 06/08/2019 | bed curtain | UM | 167 | 10 | 2 |
| DETEC-S560 | 01/09/2019 | rectal swab | U | 167 | 7 | 5 |
| DETEC-S792 | 17/09/2019 | rectal swab | FUB | 174 | 5 | 3 |
| DETEC-S565 | 01/09/2019 | rectal swab | U | 196 | 6 | 5 |
| DETEC-E601 | 03/09/2019 | sink countertop | FU | 345 | 30 | 4 |
| DETEC-P169 | 06/08/2019 | rectal swab | U | 410 | 11 | 2 |
| DETEC-P829 | 24/09/2019 | rectal swab | FU | 453 | 22 | 3 |
| DETEC-P80 | 01/08/2019 | rectal swab | U | 744 | 23 | 4 |
| DETEC-C31 | 15/10/2020 | clinical sample | FU | 1011 | 10 | 4 |
| DETEC-E1070 | 22/10/2019 | sink drain | U | 1193 | 7 | 2 |
| DETEC-P793 | 12/09/2019 | rectal swab | U | 1193 | 2 | 7 |
| DETEC-P836 | 24/09/2019 | rectal swab | U | 1193 | 7 | 2 |
| DETEC-E1005 | 15/10/2019 | sink drain | U | 1193 | 7 | 2 |
| DETEC-P881 | 08/10/2019 | oral swab | U | 1236 | 14 | 2 |
| DETEC-P649 | 05/09/2019 | rectal swab | UM | 2003 | 9 | 8 |
| DETEC-P666 | 10/09/2019 | oral swab | FU | 2003 | 2 | 12 |

[1] all complete genomes contained only circular chromosome and plasmid sequences

[2] DETEC-E = ICU environment, DETEC-P = ICU patient, DETEC-S = ICU staff, DETEC-C = clinical specimen

[3] assembly methods: U = unicycler, UM = unicycler with small plasmids finalised manually, FU = flye > unicycler, FUB = flye > unicycler bold mode, FUM = flye > unicycler with small plasmids finalised manually

[4] non-duplicate number of acquired antibiotic resistance genes

**Table S4**: Reference genomes used for phylogenetic analyses.

| ST | # isolates | Alignment reference | Alignment type | Phylogeny available |
|---|---|---|---|---|
| 1193 | 14 | MCJCHV-1 (GCF_003344465.1) | Recombination removed | Yes |
| 1236 | 5 | DETEC-P881 hybrid assembly | Recombination removed | Yes |
| 131 | 25 | EC958 (GCF_000285655.3) | Recombination removed | Yes |
| 167 | 6 | 51008369SK1 (GCF_003254065.1) | Recombination removed | Yes |
| 2003 | 6 | DETEC-P666 hybrid assembly | Recombination removed | Yes |
| 345 | 12 | DETEC-E601 hybrid assembly | Recombination removed | Yes |
| 410 | 11 | YD786 (GCF_001442495.1) | Recombination removed | Yes |
| 453 | 5 | DETEC-P829 hybrid assembly | Recombination removed | Yes |
| 744 | 8 | DETEC-P80 hybrid assembly | Recombination removed | Yes |
| 48 | 4 | DETEC-P61 hybrid assembly | Recombination removed | Yes |
| 648 | 4 | DETEC-P676 Illumina assembly | Recombination removed | Yes |
| 69 | 4 | DETEC-S586 hybrid assembly | Recombination removed | Yes |
| 10 | 3 | DETEC-S581 Illumina assembly | Full genome | No |
| 13 | 2 | DETEC-C13 Illumina assembly | Full genome | No |
| 174 | 2 | DETEC-S792 hybrid assembly | Full genome | No |
| 196 | 2 | DETEC-S565 hybrid assembly | Full genome | No |
| 354 | 2 | DETEC-P842 Illumina assembly | Full genome | No |
| 38 | 3 | DETEC-P152 Illumina assembly | Full genome | No |
| 393 | 2 | DETEC-C10 Illumina assembly | Full genome | No |
| 4456 | 2 | DETEC-C2 Illumina assembly | Full genome | No |
| 457 | 3 | DETEC-S592 Illumina assembly | Full genome | No |
| 5416 | 2 | DETEC-P817 Illumina assembly | Full genome | No |
| 58 | 2 | DETEC-P1056 hybrid assembly | Full genome | No |

**Table S5:** CTX-M genes and IS*Ecp1* transposition unit characteristics in *E. coli* isolates.

| Isolate | Genome | ST | CTX-M | TPU size | TSD | TPU location | Source of chromosomal TPU[2] |
|---|---|---|---|---|---|---|---|
| DETEC-C1 | draft | 1193 | 27 | - | - | - | - |
| DETEC-C10 | draft | 393 | 27 | - | - | - | - |
| DETEC-C11 | draft | 131 | 15 | - | - | - | - |
| DETEC-C12 | draft | 1193 | 27 | - | - | - | - |
| DETEC-C13 | draft | 13 | 55 | - | - | plasmid; X1 | - |
| DETEC-C14 | draft | 131 | 27 | 2927 | TACAA | chromosome | - |
| DETEC-C15 | draft | 131 | 14, 15 | 3060 | GCGGA | plasmid; Z | - |
| DETEC-C16 | draft | 12 | 14 | 4132 | CATTA | plasmid; FII-2 | - |
| DETEC-C17 | draft | 12742 | 55 | 3050 | ATGAA | chromosome | H12 plasmid (AP023198) |
| DETEC-C18 | draft | 457 | 65 | - | - | - | - |
| DETEC-C19 | draft | 1193 | 55 | 2971 | TACTT | plasmid; I1 | - |
| DETEC-C2 | draft | 4456 | 65 | - | - | - | - |
| DETEC-C20 | draft | 58 | 14, 55 | - | - | plasmid; FII-33 | - |
| DETEC-C21 | draft | 131 | 27 | 2991 | TTTTA | chromosome | - |
| DETEC-C22 | draft | 1193 | 3 | 2927 | TTCTT | chromosome | - |
| DETEC-C23 | draft | 131 | 14 | - | - | chromosome | - |
| DETEC-C23 | draft | 131 | 27 | - | - | - | - |
| DETEC-C24 | draft | 131 | 27 | 2991 | TTTTA | chromosome | - |
| DETEC-C25 | draft | 706 | - | - | - | - | - |
| DETEC-C26 | draft | 2179 | 65 | - | - | - | - |
| DETEC-C27 | draft | 131 | 27 | 2991 | TTTTA | chromosome | - |
| DETEC-C28 | draft | 131 | 27 | - | - | - | - |
| DETEC-C29 | draft | 131 | 27 | - | - | - | - |
| DETEC-C3 | draft | 131 | 15 | - | - | - | - |
| DETEC-C30 | draft | 410 | 15 | - | - | - | - |
| DETEC-C31 | complete | 1011 | 55 | 2971 | TACTT | plasmid; I1 | - |
| DETEC-C32 | draft | 44 | 15 | - | - | - | - |
| DETEC-C4 | draft | 131 | 14 | - | - | - | - |
| DETEC-C5 | draft | 744 | 64 | - | - | - | - |
| DETEC-C6 | draft | 453 | - | - | - | - | - |
| DETEC-C7 | draft | 1193 | 55 | 2971 | TACTT | plasmid; I1 | - |
| DETEC-C8 | draft | 4456 | 55 | 2971 | GTTTC | plasmid; FII-107:FIB-1 | - |
| DETEC-C9 | draft | 13 | 55 | - | - | plasmid; X1 | - |
| DETEC-P526 | draft | 88 | 65 | - | - | - | - |
| DETEC-E1005 | complete | 1193 | 55 | 2971 | TATAT | chromosome | - |
| DETEC-E1007 | draft | 1193 | 55 | 2971 | TATAT | chromosome | - |
| DETEC-E1010 | draft | 345 | 55 | - | - | plasmid; FII-33 | - |
| DETEC-E1024 | draft | 1193 | 55 | 2971 | TATAT | chromosome | - |
| DETEC-E1029 | draft | 648 | 14 | - | - | - | - |
| DETEC-E1033 | draft | 1236 | 55 | 2845 | TGTTT | chromosome | - |
| DETEC-E1070 | complete | 1193 | 55 | 2971 | TATAT | chromosome | - |
| DETEC-E1074 | draft | 69 | 65 | - | - | - | - |
| DETEC-E159 | draft | 48 | 64 | 18201 | TGTGT | chromosome | HI2 plasmid (MT773678) |
| DETEC-E161 | draft | 1196 | 55 | - | - | - | - |
| DETEC-E223 | complete | 167 | 14 | - | - | plasmid; FII-18 | - |
| DETEC-E232 | draft | 744 | 55 | - | - | plasmid; FII-33 | - |
| DETEC-E249 | draft | 744 | 55 | - | - | plasmid; FII-33 | - |
| DETEC-E255 | draft | 744 | 55 | - | - | plasmid; FII-33 | - |
| DETEC-E257 | draft | 744 | 55 | - | - | plasmid; FII-33 | - |
| DETEC-E387 | draft | 744 | 55 | - | - | plasmid; FII-33 | - |
| DETEC-E455 | draft | 410 | 15 | - | - | - | - |
| DETEC-E456 | draft | 410 | 15 | - | - | - | - |
| DETEC-E457 | draft | 410 | 15 | - | - | - | - |
| DETEC-E458 | draft | 410 | 15 | - | - | - | - |
| DETEC-E459 | draft | 410 | 15 | - | - | - | - |
| DETEC-E471 | draft | 167 | - | - | - | - | - |
| DETEC-E480 | complete | 131 | 14, 55 | - | - | plasmid; FII-33 | - |
| DETEC-E481 | draft | 131 | 14, 55 | - | - | plasmid; FII-33 | - |
| DETEC-E521 | draft | 1485 | 55 | 2845 | TGTTT | chromosome | - |
| DETEC-E600 | draft | 410 | 15 | - | - | - | - |
| DETEC-E601 | complete | 345 | 55, 65 | - | - | plasmid; FII-33 | - |
| DETEC-E605 | draft | 345 | 55 | - | - | plasmid; FII-33 | - |
| DETEC-E628 | draft | 131 | 14 | - | - | - | - |
| DETEC-E632 | draft | 131 | 14 | - | - | - | - |
| DETEC-E674 | draft | 131 | 14 | - | - | - | - |
| DETEC-E705 | draft | 345 | 55, 65 | - | - | plasmid; FII-33 | - |
| DETEC-E708 | draft | 345 | 65 | - | - | - | - |
| DETEC-E710 | draft | 345 | 65 | - | - | - | - |
| DETEC-E746 | draft | 2003 | 14 | 2855 | TAGTA | chromosome | - |
| DETEC-E749 | draft | 2003 | 14 | 2855 | TAGTA | chromosome | - |

| | | | | | | | |
|---|---|---|---|---|---|---|---|
| DETEC-E751 | draft | 2003 | 14 | 2855 | TAGTA | chromosome | - |
| DETEC-E799 | draft | 167 | 14 | - | - | - | - |
| DETEC-E810 | draft | 345 | 55, 65 | - | - | plasmid; FII-33 | - |
| DETEC-E812 | draft | 345 | 55, 65 | - | - | plasmid; FII-33 | - |
| DETEC-E814 | draft | 345 | 55, 65 | - | - | plasmid; FII-33 | - |
| DETEC-E816 | draft | 345 | 55, 65 | - | - | plasmid; FII-33 | - |
| DETEC-E835 | draft | 2003 | 14 | 2855 | TAGTA | chromosome | - |
| DETEC-E846 | draft | 345 | 55, 65 | - | - | plasmid; FII-33 | - |
| DETEC-E869 | draft | 617 | 55 | 5800 | TAATT | chromosome | I2 plasmid (LR890295) |
| DETEC-E880 | draft | 1193 | 55 | 2971 | TATAT | chromosome | - |
| DETEC-E882 | draft | 1236 | 55 | 2845 | TGTTT | chromosome | - |
| DETEC-E883 | draft | 1236 | 55 | 2845 | TGTTT | chromosome | - |
| DETEC-P1008 | draft | 345 | 55, 65 | - | - | plasmid; FII-33 | - |
| DETEC-P1021 | draft | 648 | 14 | - | - | - | - |
| DETEC-P1022 | draft | 5416 | 65 | - | - | - | - |
| DETEC-P1030 | draft | 1236 | 55 | 2845 | TGTTT | chromosome | - |
| DETEC-P1056 | complete | 58 | 14 | 3060 | GCGGA | plasmid; Z | - |
| DETEC-P1067 | draft | 648 | 14 | - | - | - | - |
| DETEC-P150 | draft | 162 | 14 | - | - | - | - |
| DETEC-P152 | draft | 38 | 14 | 4477 | TGAAA | chromosome | FII-2 plasmid (HM355591) |
| DETEC-P156 | draft | 4503 | 64 | - | - | chromosome | - |
| DETEC-P169 | complete | 410 | 15 | - | - | plasmid; F-type[x] | - |
| DETEC-P196 | draft | 156 | 55 | 2971 | TCATA | plasmid; HI2 | - |
| DETEC-P219 | draft | 354 | 14 | 3445 | TAACC | chromosome | I-complex plasmid (CP054459)[4] |
| DETEC-P323 | draft | 410 | 15 | - | - | - | - |
| DETEC-P351 | complete | 167 | - | - | - | - | - |
| DETEC-P353 | draft | 167 | - | - | - | - | - |
| DETEC-P363 | draft | 196 | 55 | - | - | - | - |
| DETEC-P449 | draft | 5614 | 15 | - | - | - | - |
| DETEC-P452 | draft | 410 | 15 | - | - | - | - |
| DETEC-P475 | draft | 12546 | 65 | - | - | - | - |
| DETEC-P477 | draft | 744 | 55 | - | - | plasmid; FII-33 | - |
| DETEC-P479 | draft | 453 | 55 | - | - | plasmid; FII-33 | - |
| DETEC-P530 | draft | 410 | 15 | - | - | - | - |
| DETEC-P545 | draft | 131 | 14 | - | - | - | - |
| DETEC-P546 | draft | 131 | 55 | - | - | plasmid; FII-33 | - |
| DETEC-P61 | complete | 48 | 64 | 18201 | TGTGT | chromosome | HI2 plasmid (MT773678) |
| DETEC-P622 | complete | 131 | 14, 55 | - | - | plasmid; FII-33 | - |
| DETEC-P623 | draft | 131 | 14, 55 | - | - | plasmid; FII-33 | - |
| DETEC-P649 | complete | 2003 | 14 | 2855 | TAGTA | chromosome | - |
| DETEC-P655 | draft | 69 | 65 | - | - | - | - |
| DETEC-P656 | draft | 8492 | 65 | - | - | - | - |
| DETEC-P666 | complete | 2003 | 14 | 2855 | TAGTA | chromosome | - |
| DETEC-P671 | draft | 131 | 14 | - | - | - | - |
| DETEC-P676 | draft | 648 | 14 | - | - | - | - |
| DETEC-P72 | draft | 38 | 14 | 4477 | TGAAA | chromosome | FII-2 plasmid (HM355591) |
| DETEC-P756 | draft | 3045 | 15 | 11384 | TATCA | chromosome | H12 plasmid (AP023198) |
| DETEC-P793 | complete | 1193 | 55 | 2841 | ATTCA | chromosome | - |
| | | | 55 | 3350 | ATGTT | plasmid; ColE2-like | - |
| DETEC-P794 | draft | 457 | - | - | - | - | - |
| DETEC-P80 | complete | 744 | 55 | - | - | plasmid; FII-33 | - |
| DETEC-P817 | draft | 5416 | 65 | - | - | - | - |
| DETEC-P829 | complete | 453 | 55 | 2971 | TCATA | plasmid; HI2 | - |
| DETEC-P836 | complete | 1193 | 55 | 2971 | TATAT | chromosome | - |
| DETEC-P838 | draft | 3941 | 55 | 2845 | TGTTT | chromosome | - |
| DETEC-P842 | draft | 354 | 55 | - | - | - | - |
| DETEC-P849 | draft | 569 | 65 | - | - | - | - |
| DETEC-P855 | draft | 450 | 15 | 2917 | TTTTA | chromosome | - |
| DETEC-P876 | draft | 1193 | 55 | 2971 | TATAT | chromosome | - |
| DETEC-P881 | complete | 1236 | 55 | 2845 | TGTTT | chromosome | - |
| DETEC-S525 | draft | 131 | 14, 55 | - | - | plasmid; FII-33 | - |
| DETEC-S560 | complete | 167 | 55 | 2971 | TACTT | plasmid; I1 | - |
| DETEC-S561 | draft | 1193 | 3 | 2927 | TTCTT | chromosome | - |
| DETEC-S562 | draft | 1722 | 15 | - | - | chromosome | - |
| DETEC-S563 | draft | 453 | 55 | - | - | - | - |
| DETEC-S564 | draft | 871 | 55 | - | - | plasmid; X1 | - |
| DETEC-S565 | complete | 196 | 55 | 2971 | TACTT | plasmid; I1 | - |
| DETEC-S566 | complete | 131 | 14 | 3060 | GCGGA | plasmid; Z | - |
| DETEC-S567 | draft | 3052 | 15 | 11384 | TAGCA | chromosome | H12 plasmid (AP023198) |
| DETEC-S568 | draft | 48 | 55 | - | - | - | - |
| DETEC-S569 | draft | 117 | 55 | - | - | - | - |
| DETEC-S570 | draft | 393 | 27 | - | - | - | - |

| | | | | | | | |
|---|---|---|---|---|---|---|---|
| DETEC-S571 | draft | 69 | 14 | 3242 | TATTA | chromosome | - |
| DETEC-S572 | draft | 202 | 15 | - | - | - | - |
| DETEC-S573 | draft | 131 | 55 | 6397 | TTGAT | chromosome | I1 plasmid (pDETEC69/pDETEC73) |
| DETEC-S575 | draft | 708 | 15 | - | - | - | - |
| DETEC-S576 | draft | 5700 | 27 | - | - | - | - |
| DETEC-S577 | draft | 453 | 55 | - | - | - | - |
| DETEC-S578 | draft | 48 | 55 | - | - | - | - |
| DETEC-S579 | draft | 2914 | 27 | - | - | - | - |
| DETEC-S580 | draft | 38 | 14 | - | - | chromosome | - |
| DETEC-S581 | draft | 10 | 14 | - | - | - | - |
| DETEC-S582 | draft | 174 | 15 | - | - | - | - |
| DETEC-S583 | draft | 624 | 14 | - | - | - | - |
| DETEC-S584 | draft | 2690 | 55 | - | - | - | - |
| DETEC-S585 | draft | 1249 | 14 | - | - | - | - |
| DETEC-S586 | complete | 69 | 24 | 3271 | AATCA | plasmid; X4 | - |
| DETEC-S587 | draft | 1163 | 27 | - | - | - | - |
| DETEC-S589 | complete | 95 | 14 | 3060 | GCGGA | plasmid; Z | - |
| DETEC-S590 | draft | 10 | 55 | - | - | plasmid; X1 | - |
| DETEC-S591 | draft | 773 | 14 | 3502 | TGATA | chromosome | - |
| DETEC-S592 | draft | 457 | 27 | - | - | - | - |
| DETEC-S790 | draft | 1968 | 55 | - | - | plasmid; FII-33 | - |
| DETEC-S791 | draft | 93 | 55, 65 | 2971 | TACTT | plasmid; I1 | - |
| DETEC-S792 | complete | 174 | 55 | - | - | plasmid; FII-2 | - |
| DETEC-S897 | draft | 10 | 14 | - | - | plasmid; Z | - |

[1] - = not present or not determined.

[2] plasmid type is listed, along with a GenBank accession (in brackets) for a representative of the lineage that is the likely source of the chromosomal TPU.

[3] sub-type: FII-31:FII-36:FIA-4:FIB-58.

[4] the plasmid at accession CP054459 does not contain an IS*Ecp*1-*bla*$_{CTX-M}$ TPU, and represents an ancestral form of the plasmid that was the likely the source of this chromosomal TPU.

**Table S6:** Diversity amongst *E. coli* isolates of the same ST.

| ST | Total isolates | Patient isolates | Environmental isolates | Staff isolates | Clinical isolates | Distinct people isolated from | Maximum SNPs between isolates of this ST |
|---|---|---|---|---|---|---|---|
| 131 | 25 | 5 | 5 | 3 | 12 | 6 | 597 |
| 1193 | 14 | 3 | 5 | 1 | 5 | 5 | 136 |
| 345 | 12 | 2 | 10 | - | - | 1 | 99 |
| 410 | 11 | 4 | 6 | - | 1 | 2 | 212 |
| 744 | 8 | 2 | 5 | - | 1 | 4 | 130 |
| 167 | 6 | 2 | 3 | 1 | - | 4 | 272 |
| 2003 | 6 | 2 | 4 | - | - | 2 | 8 |
| 1236 | 5 | 2 | 3 | - | - | 1 | 1 |
| 453 | 5 | 2 | | 2 | 1 | 5 | 557 |
| 48 | 4 | 1 | 1 | 2 | - | 3 | 1504 |
| 648 | 4 | 3 | 1 | - | - | 2 | 178 |
| 69 | 4 | 1 | 1 | 2 | - | 4 | 311 |
| 10 | 3 | - | - | 3 | - | 3 | 20765 |
| 38 | 3 | 2 | - | 1 | - | 2 | 9016 |
| 457 | 3 | 1 | - | 1 | 1 | 3 | 7299 |
| 13 | 2 | - | - | - | 2 | 1 | 3 |
| 174 | 2 | - | - | 2 | - | 2 | 1 |
| 196 | 2 | 1 | - | 1 | - | 2 | 9949 |
| 354 | 2 | 2 | - | - | - | 2 | 9132 |
| 393 | 2 | - | - | 1 | 1 | 2 | 336 |
| 4456 | 2 | - | - | - | 2 | 1 | 1774 |
| 5416 | 2 | 2 | - | - | - | 2 | 1 |
| 58 | 2 | 1 | - | - | 1 | 2 | 8829 |

**Table S7:** Maximum cgSNPs amongst *E. coli* isolates of the same ST associated with a single patient

| Patient ID | ST | # SNPs |
|---|---|---|
| P1 | 38 | 0 |
| P5 | 48 | 0 |
| P3 | 131 | 9 |
| P14 | 131 | 3 |
| P11 | 167 | 12 |
| P8 | 345 | 99 |
| P6 | 410 | 4 |
| P23 | 648 | 12 |
| P3 | 744 | 3 |
| P23 | 1193 | 8 |
| P27 | 1193 | 0 |
| P24 | 1236 | 1 |
| P17 | 2003 | 1 |

**Table S8:** GenBank plasmids containing the I1 plasmid TACTT-flanked IS*Ecp1*-*bla*$_{CTX-M-55}$ TPU

| Plasmid | Accession | Host | Country | Year | Source | Size (bp) | *repA*[1] |
|---|---|---|---|---|---|---|---|
| p6607-69 | CP045527 | *S. sonnei* | Switzerland | 2017 | human faeces | 89,848 | pDETEC69 |
| pST53-2 | CP050747 | *S. enterica* Typhimurium | China: Shanghai | 2011 | human faeces | 88,006 | pDETEC69 |
| pHNRD174 | KX246268 | *E. coli* | China: Guangdong | ≤2020 | duck | 86,207 | pDETEC69 |
| pXH990_3 | CP019358 | *E. coli* | China: Zhejiang | 2016 | human urine | 86,028 | pDETEC69 |
| pD3-B | CP010142 | *E. coli* | China: Jiangsu | 2014 | dog | 90,074 | pDETEC69 |
| p628-CTXM | KP987217 | *K. pneumoniae* | China: Beijing | 2010 | human cerebrospinal fluid | 85,338 | pDETEC69 |
| pSKLX3330 | KJ866866 | *E. coli* | China: Zhejiang | ≤2020 | human urine | 89,672 | pDETEC69 |
| pKP4823_3 | KF790923 | *K. pneumoniae* | China: Zhejiang | 2019 | human urine | 86,182 | pDETEC69 |
| pURN1-2021 | CP082826 | *E. coli* | Kazakhstan | 2021 | human urine | 86,164 | pDETEC69 |
| p14523A | CP074429 | *Salmonella* sp. | China: Xinjiang | 2016 | human urine | 85,862 | pDETEC69 |
| pRHBSTW-00218_2 | CP056650 | *E. hormachei* | United Kingdom | 2017 | wastewater influent | 86,191 | pDETEC69 |
| pWP3-S18-ESBL-09 | AP022037 | *E. coli* | Japan | 2018 | wastewater effluent | 88,528 | pDETEC69 |
| pKP4823_3 | CP082793 | *K. pneumoniae* | China: Zhejiang | 2019 | human urine | 86,182 | pDETEC69 |
| pEC32-IncI1 | CP085621 | *E. coli* | China: Guangdong | 2014 | human urine | 88,542 | pDETEC69 |
| pS29-IncI1 | CP085700 | *S. enterica* | China: Guangdong | 2014 | human faeces | 88,502 | pDETEC69 |
| pKko009_2 | CP091675 | *E. coli* | Japan | 2012 | river water | 89,404 | pDETEC69 |
| pEC24-3 | CP060887 | *E. coli* | China: Zhejiang | 2017 | human urine | 86,975 | pDETEC69 |
| pEC20-3 | CP060904 | *E. coli* | China: Zhejiang | 2017 | human throat swab | 86,975 | pDETEC69 |
| pEC19-3 | CP060910 | *E. coli* | China: Zhejiang | 2017 | human urine | 86,975 | pDETEC69 |
| p2474-3 | CP021208 | *E. coli* | China: Hefei | 2015 | human blood | 86,725 | pDETEC73 |
| pKFu019_1 | CP091698 | *E. coli* | Japan | 2011 | river water | 86,740 | pDETEC73 |
| pEC7-3 | CP060965 | *E. coli* | China: Zhejiang | 2016 | human urine | 89,379 | pDETEC73 |
| p2-3 | CP091573 | *S. enterica* | China: Zhejiang | 2016 | human faeces | 86,718 | pDETEC73 |
| p20 | CP099778 | *S. sonnei* | Belgium | 2015 | human | 86,718 | pDETEC73 |
| p72_2 | CP101556 | *K. pneumoniae* | China: Yunnan | 2013 | human urine | 87,511 | pDETEC73 |

[1] Indicates whether the plasmid contains the *repA* variant found in the pDETEC69 or pDETEC73 backbone.

**Table S9:** GenBank plasmids containing the Z plasmid GCGGA-flanked IS*Ecp1-bla*$_{CTX-M-14}$ TPU

| Plasmid | Accession | Host | Country[1] | Year | Source | Size (bp) |
|---|---|---|---|---|---|---|
| pCT[2] | FN868832 | *E. coli* | UK | 2004 | dairy farm calves | 93,629 |
| p2D-CTX-M-14 | CP059004 | *E. coli* | China | 2019 | human blood | 96,449 |
| pWP7-S17-ESBL-01_1 | AP022174 | *E. coli* | Japan | 2017 | wastewater plant effluent | 94,776 |
| pWP8-S17-ESBL-12_2 | AP022224 | *E. coli* | Japan | 2017 | wastewater plant effluent | 87,171 |
| pBEC1-S17-ESBL-07_1 | AP022296 | *E. coli* | Japan | 2017 | oceanic water | 105,884 |
| unnamed | LR890721 | *K. pneumoniae* | Australia* | - | - | 89,062 |
| pJX1-2 | CP064254 | *K. pneumoniae* | China | 2018 | human | 102,456 |
| pF16EC0557-1 | CP088388 | *E. coli* | South Korea | 2016 | human blood | 150,230 |
| pF16EC0617-4 | CP088378 | *E. coli* | South Korea | 2016 | human blood | 87,357 |
| pF16EC0342-2 | CP088412 | *E. coli* | South Korea | 2016 | human blood | 111,606 |
| pF16EC0121-1 | CP088449 | *E. coli* | South Korea | 2016 | human blood | 94,534 |
| pE16EC0790-2 | CP088518 | *E. coli* | South Korea | 2016 | human blood | 90,665 |
| pC17EC0264-2 | CP088619 | *E. coli* | South Korea | 2017 | human blood | 94,108 |
| pD16EC1060-3 | CP088610 | *E. coli* | South Korea | 2016 | human blood | 98,480 |
| pB16EC1060-3 | CP088732 | *E. coli* | South Korea | 2016 | human blood | 90,665 |
| pB16EC0725-1 | CP088777 | *E. coli* | South Korea | 2016 | human blood | 106,218 |
| pB16EC0268-1 | CP088807 | *E. coli* | South Korea | 2016 | human blood | 90,653 |
| pA17EC0191-3 | CP088824 | *E. coli* | South Korea | 2017 | human blood | 93,421 |
| pA16EC0054-1 | CP088870 | *E. coli* | South Korea | 2016 | human blood | 88,005 |
| unnamed3 | CP090250 | *E. coli* | China | 2018 | human | 90,981 |
| p3 | CP090538 | *S. enterica* | China | 2016 | human | 90,380 |
| pEC-16-35-4 | CP093229 | *E. coli* | China | 2016 | human ascites | 90,339 |
| pEC9682-2 | CP095273 | *E. coli* | China | 2021 | human vaginal secretion | 96,608 |
| pPIB-2 | CP090404 | *Shigella* sp. | China | 2014 | human | 87,994 |
| P2 | OX030730 | *E. coli* | Spain | - | - | 100,782 |
| p69 | CP099769 | *S. sonnei* | Belgium | 2018 | human | 111,597 |
| pSH15sh99 | KY471628 | *S. sonnei* | China | 2015 | drinking water | 104,285 |
| pSH15sh104 | KY471629 | *S. sonnei* | China | 2015 | human faeces | 104,285 |
| RCS56_p | LT985270 | *E. coli* | France* | - | - | 90,206 |
| RCS68_p | LT985278 | *E. coli* | France* | - | - | 86,943 |
| pSH262-2 | MG299128 | *S. sonnei* | China | 2016 | human | 109,845 |
| pSH271-2 | MG299131 | *S. sonnei* | China | 2016 | water | 109,845 |
| pSH272-2 | MG299133 | *S. sonnei* | China | 2016 | human | 109,845 |
| pSH284-2 | MG299147 | *S. sonnei* | China | 2016 | human | 109,845 |
| pSH287-2 | MG299151 | *S. sonnei* | China | 2016 | human | 109,845 |
| p50579417_2 | CP033883 | *E. coli* | Norway | 2012 | human urine | 88,005 |
| unnamed | LR595871 | *E. coli* | UK* | - | human faeces | 94,061 |
| unnamed | LR595872 | *E. coli* | UK* | - | human faeces | 96,306 |
| unnamed | LR595877 | *E. coli* | UK* | - | human faeces | 111,594 |
| unnamed | LR595880 | *E. coli* | UK* | - | human faeces | 96,305 |
| unnamed | LR595888 | *E. coli* | UK* | - | human faeces | 96,306 |
| unnamed | LR595889 | *E. coli* | UK* | - | human faeces | 94,296 |
| pKP16-19-tet(A) | MN480462 | *K. pneumoniae* | China | 2016 | human sputum | 106,623 |
| pSCU-103-3 | CP054460 | *E. coli* | USA | 2015 | human rectal swab | 34,914 |

[1] * indicates where the submitting authors' location is listed, as a country of isolation was not stated in the GenBank entry.

[2] pCT is has a K-type replicon, indicating that recombination has likely resulted in the movement of the GCGGA-flanked IS*Ecp1-bla*$_{CTX-M-14}$ TPU and adjacent backbone sequence between plasmids with Z and K-type replicons.

**Table S10:** Complete plasmid sequences generated in this study.

| **Type**<br>Sub-type | Plasmid | Size (bp) | #ARGs |
|---|---|---|---|
| **F-type** | | | |
| FII-2 | pDETEC44 | 82,435 | 5 |
| FII-16 | pDETEC24 | 76,416 | 3 |
| FII-29 | pDETEC80 | 81,583 | 2 |
| FII-33 | pDETEC21 | 103,125 | 8 |
| FII-33 | pDETEC61 | 93,520 | 8 |
| FII-33:N | pDETEC56 | 103,383 | 8 |
| FII-33:N | pDETEC60 | 82,763 | 3 |
| FII-35 | pDETEC48 | 70,464 | - |
| FII-49 | pDETEC57 | 13,744 | 1 |
| FIB-54:phage-plasmid | pDETEC37 | 142,064 | 9 |
| FII-1:FIA-1 | pDETEC32 | 98,478 | 1 |
| FII-36:FIA-4 | pDETEC5 | 116,501 | 5 |
| FII-18:FIB-1 | pDETEC11 | 154,454 | 8 |
| FII-18:FIB-1 | pDETEC19 | 159,599 | 7 |
| FII-18:FIB-1 | pDETEC68 | 102,896 | 4 |
| FII-18:FIB-1 | pDETEC77 | 132,207 | 2 |
| FII-18:FIB-1 | pDETEC88 | 111,058 | 2 |
| FII-18:FIB-35 | pDETEC81 | 90,243 | - |
| FII-107:FIB-1 | pDETEC46 | 141,890 | 4 |
| FIA-1:FIB-10 | pDETEC3 | 108,072 | 6 |
| FIA-1:FIB-10 | pDETEC15 | 82,761 | - |
| FIA-6:FIB-20 | pDETEC55 | 107,932 | 5 |
| FII-2:FIA-1:FIB-1 | pDETEC38 | 128,466 | 3 |
| FII-18:FIA-5:FIB-1 | pDETEC23 | 178,682 | 6 |
| FII-18:FIA-5:FIB-1 | pDETEC72 | 187,674 | 5 |
| FII-18:FIA-2:FIB-8 | pDETEC66 | 194,772 | - |
| FII-novel:FIA-1:FIB-23 | pDETEC25 | 151,807 | 8 |
| FII-31:FII-36:FIA-4:FIB-58 | pDETEC2 | 113,066 | 6 |
| **I-complex** | | | |
| I1 | pDETEC6 | 77,960 | 1 |
| I1 | pDETEC14 | 87,921 | - |
| I1 | pDETEC20 | 118,716 | 4 |
| Z | pDETEC33 | 86,916 | - |
| Z | pDETEC43 | 98,708 | - |
| Z | pDETEC45 | 81,965 | - |
| I1 | pDETEC47 | 87,625 | - |
| I1 | pDETEC67 | 118,588 | 5 |
| I1 | pDETEC69 | 86,167 | 1 |
| I1 | pDETEC73 | 90,981 | 1 |
| Z | pDETEC78 | 89,022 | 1 |
| Z | pDETEC79 | 87,662 | 1 |
| Z | pDETEC82 | 88,352 | 1 |
| I1 | pDETEC89 | 86,715 | 1 |
| I1 | pDETEC91 | 85,862 | 1 |
| **X-type** | | | |
| | pDETEC35 | 38,312 | - |
| | pDETEC36 | 30,896 | - |
| X1 | pDETEC39 | 44,960 | 7 |
| X4 | pDETEC49 | 42,640 | 1 |
| | pDETEC62 | 36,682 | - |
| **H-type** | | | |
| | pDETEC13 | 209,927 | - |
| HI2 | pDETEC65 | 266,247 | 17 |

| | | | |
|---|---|---|---|
| **N-type** | | | |
| N2 | pDETEC50 | 41,594 | - |
| **phage-plasmid** | | | |
| | pDETEC1 | 119,948 | 5 |
| | pDETEC34 | 47,568 | - |
| | pDETEC74 | 89,020 | - |
| | pDETEC87 | 113,249 | - |
| **θ-RNA** | | | |
| | pDETEC9 | 3,684 | - |
| | pDETEC10 | 3,174 | - |
| | pDETEC12 | 6,200 | 2 |
| | pDETEC17 | 5,631 | - |
| | pDETEC26 | 5,775 | - |
| | pDETEC28 | 4,593 | - |
| | pDETEC40 | 5,903 | - |
| | pDETEC42 | 3,263 | - |
| | pDETEC51 | 7,939 | - |
| | pDETEC53 | 4,715 | - |
| | pDETEC63 | 4,045 | - |
| | pDETEC64 | 3,731 | - |
| | pDETEC71 | 2,678 | - |
| | pDETEC75 | 6,647 | - |
| | pDETEC83 | 7,559 | - |
| | pDETEC90 | 3,537 | - |
| **θ-Rep** | | | |
| | pDETEC7 | 6,074 | - |
| | pDETEC8 | 4,073 | - |
| | pDETEC16 | 8,520 | 1 |
| | pDETEC18 | 4,060 | - |
| | pDETEC22 | 4,692 | - |
| | pDETEC27 | 5,167 | - |
| | pDETEC30 | 4,069 | - |
| | pDETEC41 | 4,867 | - |
| | pDETEC52 | 5,163 | - |
| | pDETEC54 | 4,059 | - |
| | pDETEC58 | 4,087 | - |
| | pDETEC84 | 5,728 | - |
| | pDETEC85 | 4,296 | - |
| **Rolling-circle** | | | |
| | pDETEC4 | 2,101 | - |
| | pDETEC29 | 4,396 | - |
| | pDETEC31 | 1,551 | - |
| | pDETEC59 | 1,552 | - |
| | pDETEC76 | 1,565 | - |
| **Untyped** | | | |
| | pDETEC70 | 4,036 | - |
| | pDETEC86 | 1,985 | - |

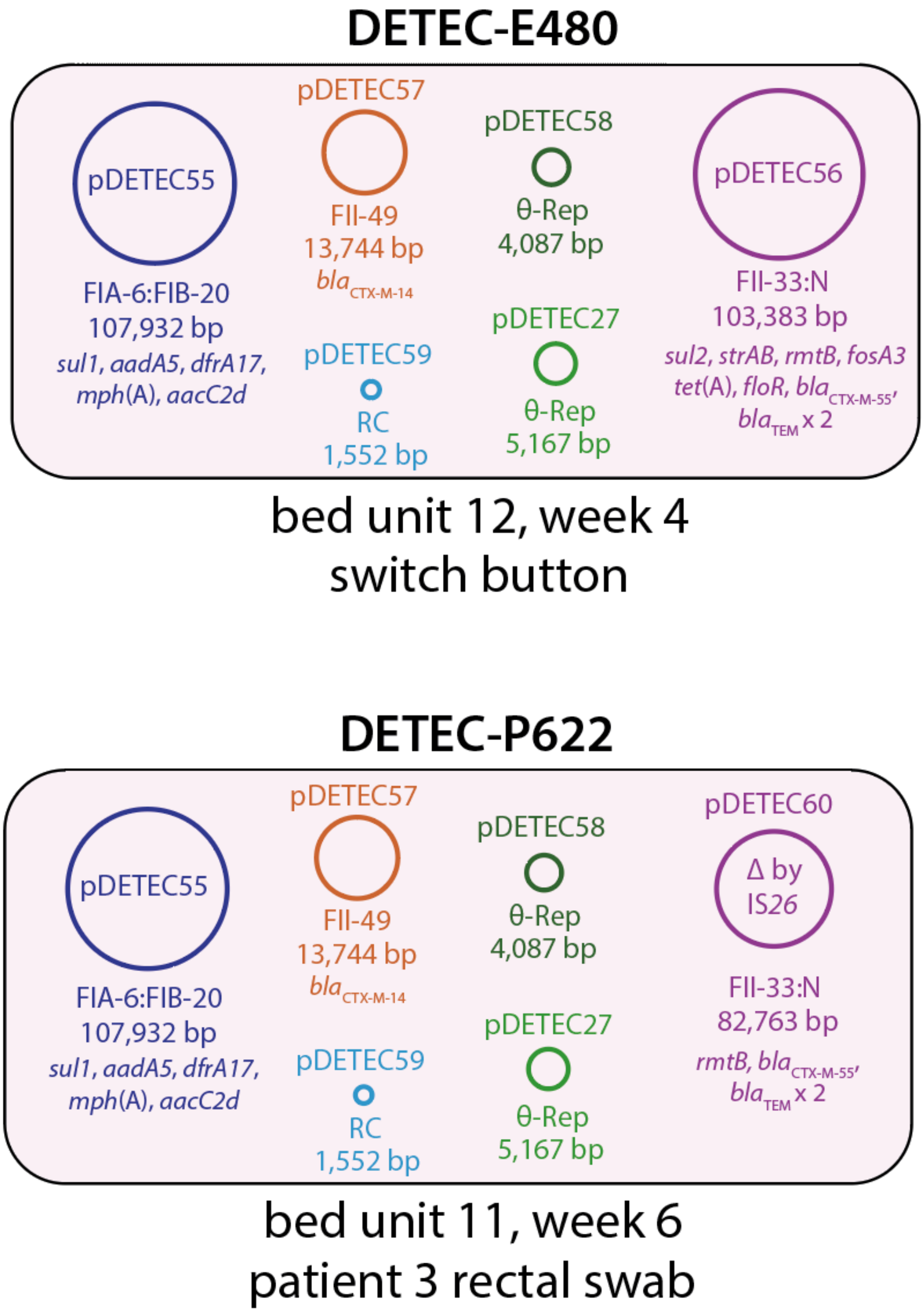


**Figure S1:** Plasmid content of ST131 isolates DETEC-E480 and DETEC-P622.
